# Developmental Characterization of Neuronal Migration Anomalies and Axon Proliferation in mTOR pathway-associated Malformations of Cortical Development

**DOI:** 10.1101/2023.03.11.532231

**Authors:** Paige Hoffman, Matthew N. Svalina, Chiara Flores, Christine Brzezinski, J. Keenan Kushner, Brandon Staple, Santos Franco, Allyson L. Alexander

## Abstract

Drug-resistant epilepsy (DRE) is a prevalent problem in children that can lead to abnormal development and various psychiatric comorbidities. Malformations of cortical development (MCD) include focal cortical dysplasia, tuberous sclerosis complex and hemimegalencephaly, which are the most common pathologies among children who undergo surgical resection for treatment of DRE. These disorders share many histopathological features, including dyslamination of the cerebral cortex and enlarged neuronal somata. Recently, genetic mutations in the mammalian target of rapamycin (mTOR) signaling cascade have been shown to underpin most MCDs. Rodent models, including the Rheb^CA^ model, recapitulate histologic and physiologic aspects of human DRE. However, there have been few studies characterizing the developmental time point of the histological changes seen in MCDs. In this study, we use *in utero* electroporation to upregulate the Rheb protein (directly upstream of mTOR) in a focal area of the neocortex. We demonstrate that mTOR dysregulation leads to focal dyslamination and increased neuronal size that is histologically similar to MCD, which correlates to spontaneous recurrent seizures. We used immunohistochemistry to investigate neuronal lamination at several time points during development between E18 and P21 and show early differences in lamination that persisted through development. Furthermore, the increased axonal length associated with mTOR upregulation occurs early in development. Our study provides a time frame for the initial development of abnormal neuronal migration and cellular growth that occurs in MCDs, and our data supports that these anatomical changes may contribute to the formation of epileptic networks.

## INTRODUCTION

Childhood epilepsy is a common disorder with a reported prevalence ranging from 0.5%-4.4%^1-9^. Uncontrolled childhood epilepsy can lead to devastating consequences including a fourteen-fold increase in premature mortality^10^. Patients are also at risk for decreased quality of life, developmental delay, conduct disorders, depression and anxiety^4,11-14^. Approximately 30-40% of patients with active epilepsy develop drug-resistant epilepsy (DRE). The most common pathologies leading to surgically-treated DRE in the pediatric population are malformations of cortical development (MCDs)^15,16^. These disorders include focal cortical dysplasia (FCD), tuberous sclerosis complex (TSC), and hemimegalencephaly (HMG). These congenital abnormalities are the most common cause of surgically-treated epilepsy in pediatric patients who have failed medical management.

Histopathological analysis of MCDs reveals abnormal migration, proliferation, and lamination of cortical neurons as well as enlarged neuronal somata^17-25^. mTOR is a master regulator of transcription and translation, promotes cellular growth, and regulates intracellular autophagy^26-28^. A growing body of evidence has demonstrated pathological mutations in the mammalian target of rapamycin (mTOR) signaling pathway in a majority of MCDs including germline mutations in patients with TSC and somatic mutations in patients with FCD and HMG^7,29-66^. These studies have demonstrated mutations in many genes within the mTOR pathway, including the mTOR activator RHEB (Fig. 1A)^39,46^. Based on the genetics of MCDs, animal models have been created to study this pathology, specifically using the constitutively active form of Rheb (Rheb^CA^) in which *in utero* electroporation is used to focally upregulate mTOR. The Rheb^CA^ mice have an area of cortex that demonstrates dyslamination and enlarged somata and is thus very similar to an area of FCD or a cortical tuber of TSC. Furthermore, these animals develop spontaneous seizures^67^.

**Figure 1.**
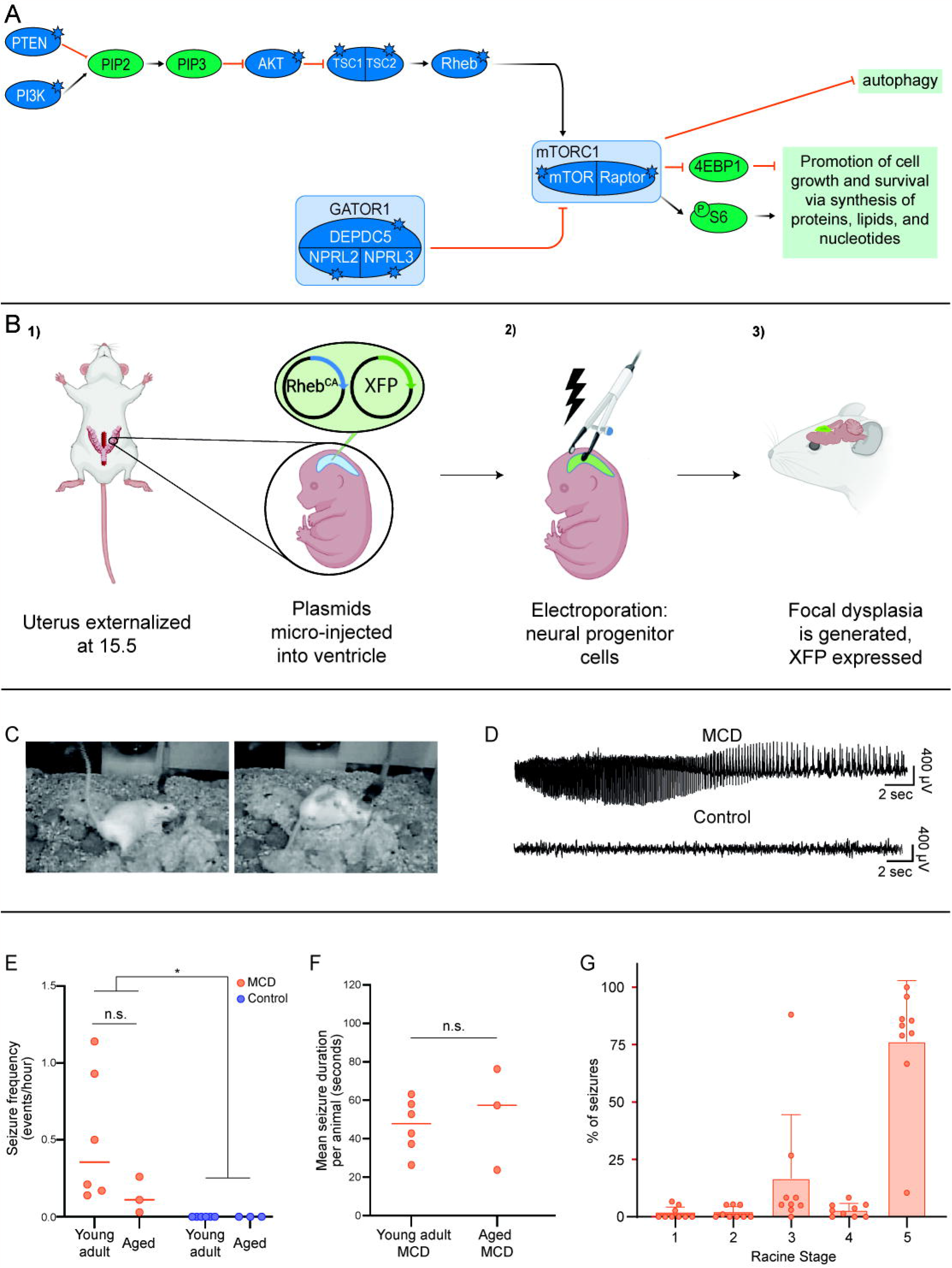
Video-Electrocorticography (EEG) recordings and demonstrate spontaneous seizures in a murine model of MCD. **(A)** Simplified mTOR activation cascade. Mutations in patients with MCDs have been shown in all proteins represented by blue starred circles. **(B)** Schematic of FCD mouse model via in utero electroporation. Note that control animals receive the XFP plasmid only, while the MCD animals receive Rheb^CA^ + XFP. **(C)** Still images from video-EEG recording in an MCD mouse demonstrates a Racine stage 5 seizure: hunching with tail up (left), losing balance and falling over (right). **(D, top)** EEG activity from left hippocampal electrode in the MCD mouse during the stage 5 seizure shown in (C) demonstrates clear ictal activity. **(D, bottom)** EEG activity from the left hippocampal electrode of a control mouse during a quiet awake period. **(E)** Seizure frequency is presented for young adult (4-6 weeks) and aged (25-26 weeks) MCD and control mice. There were no seizures recorded from any of the control mice. Bars at median per group. **(F)** Mean seizure duration (sec) per animal compared between young adult and aged MCD mice. Control mice not presented as none had seizures. Bars at median per group. **(G)** Percentage of seizures identified in MCD mice at every stage on the Racine scale. Error bars set at mean with standard deviation.

MCD is a disorder of development that has been hypothesized to be secondary to a somatic mutation during early embryogenesis in which a selected population of neurons receive the “hit” and this is propagated via clonal expansion^42,68^. To date, there have been few studies of the abnormal development of neuronal migration during development in these animal models. However, it has been shown that mTOR-associated growth of somata and dendrites is increased between birth and the third week of postnatal development in a mouse model^69^. Additionally, one study demonstrates that migration through the cortical plate is affected early in development in a model where electroporation is done early at E13.5^70^.

In this study, we used the Rheb^CA^ model of MCD to examine histological changes seen in this disorder during a developmental time course, looking at 5 developmental time points between E18 and P21. We verified that the animals in this model demonstrate spontaneous recurrent seizures. We show that histopathological changes of MCD, including migration defects and remarkable axon growth occur by postnatal day 2. We also demonstrate the surprising finding that some non-electroporated neurons in MCD animals exhibit upregulated mTOR signaling.

## METHODS

### Animals

All animal experiments were approved by the University of Colorado Anschutz Institutional Care and Use Committee. Every effort was made to minimize the number of animals. Analgesia and euthanasia methods were chosen under the advisement of the veterinarians at our institution.

### In utero electroporation

In utero electroporation was performed at E15.5 in CD1 timed-pregnant dams (Charles River, Wilmington, MA) under general anesthesia with a ECM830 ElectroSquarePorator (Harvard Apparatus, Holliston, MA). MCD mice were electroporated with the Rheb^CA^ plasmid ^67,71^ (a kind gift from Dr. Maehama^72^) and either GFP or BFP as a fluorescent marker. Control mice were electroporated with TdTomato or BFP alone. (GFP, BFP, TdTomato collectively referred to as XFP). Bupivacaine along with either carprofen or meloxicam were used as analgesics post-operatively (MWI, Boise, ID). Once born, pups were checked for fluorescent expression through the skull using a Nightsea Dual Fluorescent Protein (DFP) flashlight.

### Immunohistochemistry

Mice above the age of P0 were injected in the lower abdomen with Fatal Plus (Vortech Pharmaceuticals Ltd.) at a ratio of 2.6 uL/g body weight then allowed to rest until fully anesthetized. Perfusions were completed with phosphate buffered saline (PBS, VWR) to clear and then with 4% paraformaldehyde (PFA, VWR). Brains were removed from the skull and fixed overnight in 4% PFA. Sectioning was performed on a Compresstome 300-0Z (Precisionary Instruments, Vortich MA) at 30-50 μm. dissected from the skull and placed directly into well plates filled with 4% PFA overnight before being gently washed with room temperature PBS and sectioned. On all embryonic mice, the dam was euthanized using cervical dislocation and pups were collected from the uterine horn. In place of perfusion, embryonic pups were euthanized via hypothermia. A block of dry ice was covered by an absorbent pad to prevent freezer burn on the skin, then pups were allowed to rest until body temperatures dropped to fatal levels. Following this, all pups were decapitated and heads were placed directly into well plates filled with 4% PFA for 2 days before being washed, dissected, and sectioned.

### EEG analysis

Electrodes were implanted bilaterally into the hippocampus and cortex of mice at either 4-6 weeks of age or 5-6 months of age, under general anesthesia. The location of the cortical electrodes was chosen to correspond to the fluorescent area of MCD. All electrodes were implanted by the In Vivo Neurophysiology EEG Core on the Anschutz Medical Campus. The corresponding data files for each mouse were imported into. Analysis was performed, blinded to experimental group, using Pinnacle Technologies’ Sirenia Seizure software volume 2.0.4. Both male and female animals were used. Seizures were defined as abnormal electrical activity with corresponding behavioral changes lasting for lasted for at least 10 seconds. The Racine Scale was utilized to determine seizure severity as follows: Stage 1: freezing; Stage 2: facial twitch, automatisms, head nodding; Stage 3: forelimb clonus; Stage 4: forelimb or hindlimb clonus with rearing; Stage 5: losing balance, falling over, tail up^73^. After recording, mice were euthanized and brains were dissected for immunohistochemistry. Electrode locations were verified in a subset of animals using cresyl violet stains (data not shown).

### Cell counting

All cell counts took place via Neurolucida using contour tracing and placed markers to specify cell types. Contours were created along the central-most section of the electroporated region within a slide, and adjusted to keep cell counts between 180 to 200 for layer markers or 950 to 1080 for mTOR upregulation markers. Cells along the contour line were only counted if over half the cell was confirmed to be touching and/or within the boundary. All counts were completed on images using the closest artificial color from the emission range specified by the secondary antibody (i.e. Dk-Ms 650 used microscope filter Cy5 and cells were colored white for counting). Secondary contour borders were placed along the pia and white matter as reference points to help calculate marker distance along the contour. All counts that were completed over quarantine via Remote PC access were performed in the same dark-room setting as counts on the microscope machinery to control for variability in light and clarity. Analysis was conducted using Neurolucida Explorer’s Marker and Contour Analysis feature.

### Statistics

Statistical analysis was performed in GraphPad Prism 9.2 with significance set at α =0.05. Normality was assessed using the D’Agostino-Pearson test. For two-way ANOVA analyses, posthoc multiple comparisons were run with Tukey’s multiple comparisons tests. Kolmogorov-Smirnov tests were used to compare cumulative probability graphs and Multiple Mann-Whitney tests, using the two-stage step-up method for false discovery rates to correct for multiple comparisons, with the Desired FDR set at 5%, were used to evaluate cell distributions between groups for all ages.

## RESULTS

### Validation of Rheb^CA^ model of MCD-related epilepsy

We chose to use a previously described histologically and genetically accurate mouse model of MCD^67,71^. This model is based on the upregulation of Rheb, a GTPase that upregulates mTORC1^74^ (Fig. 1A). We created the MCD by electroporation of the constitutively active Rheb plasmid (Rheb S16H, Rheb^CA 72^) at E15.5 into the lateral ventricle (Fig. 1B). At this embryonic age, neuronal precursor cells are destined to become layer II/III pyramidal neurons. Video-Electroencephalography (EEG) was performed in 9 MCD and 9 control mice at both young adult (4-6 weeks) and aged (5-6 months) time points. All MCD mice (n = 6 young adult, n = 3 aged) and no control mice (n = 6 young adult, n = 3 age) displayed behavioral (Fig. 1C) and electrographic (Fig. 1D) seizures. There was a higher seizure frequency in mutant vs. control animals overall but only in the young adult animals on posthoc analysis (Two-way ANOVA: effect of genotype, p = 0.025, young adult mutant vs. young adult control, p = 0.019, aged mutant vs aged control, p = 0.9, Fig. 1D). There was a trend toward lower seizure frequency in the aged mice, but this was not significant (Mutant aged vs. mutant young adult, p = 0.2). There was no difference in seizure duration between young adult and aged MCD mice (median ± SEM: young adult = 47.81±5.63s, aged = 57.42±15.3s, p = 0.9049, Mann-Whitney, Fig. 1F). The most common seizure type was Racine stage 5 (Fig. 1E). Thus, we have validated that in the Rheb^CA^ mouse model of MCD the mice exhibit spontaneous recurrent seizures.

### Most electroporated cells in the Rheb^CA^ MCD model are neurons

As a further validation of the model, we quantified the number of electroporated cells which were neurons, as evidenced by the expression of NeuN, a protein that is expressed only in the nuclei of post-mitotic neurons^75^. We performed immunohistochemistry to evaluate the proportion of EP cells that co-expressed NeuN at four time points during postnatal development: P2, P7, P14 and P21 (Fig. 2A). Overall, the majority of EP neurons in both groups were also positive for NeuN (all ages combined: MCD: 95.3±1.6%, CON: 87.8±3.9%). Interestingly, statistical analysis demonstrated that significantly more EP calls are NeuN positive than control cells (two-way ANOVA, effect of genotype, p = 0.0001). The proportion of NeuN positive cells did not change significantly during development (P2: MCD =93.1 ± 2.9% ; CON = 85.3 ± 2.8%; P7: MCD =95.8 ± 1.7%; CON = 93.5 ± 1.3%; P14: MCD =95.8 ± 2.7%; CON = 85.2 ± 2.9%). Given that neurogenesis is complete before P2, there was no significant effect of age on the expression of NeuN (two-way ANOVA, p = 0.1879). This result, that more control EP cells then MCD EP cells express a neuronal phenotype, was unexpected. We postulate

**Figure 2.**
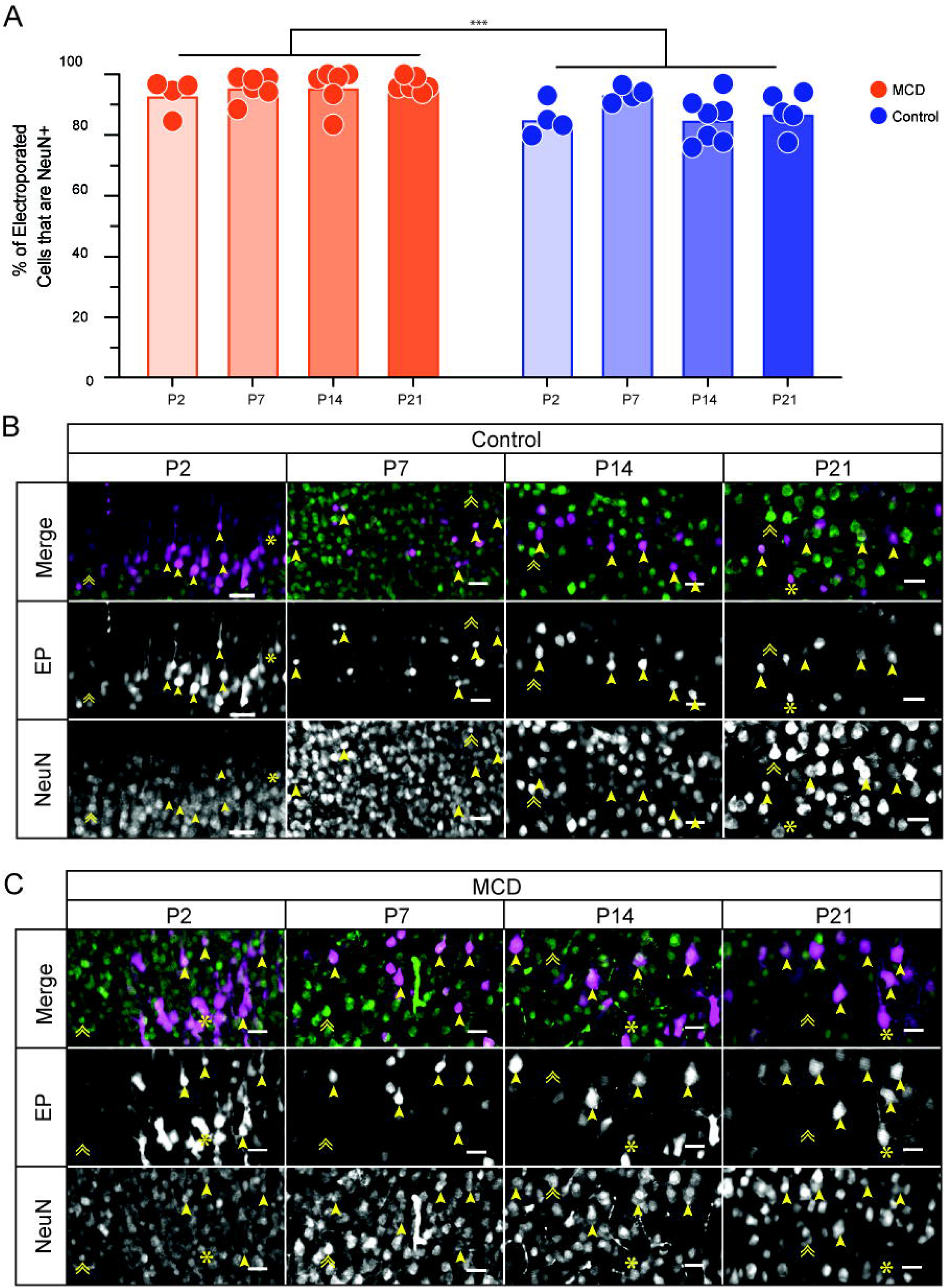
Most electroporated cells are neurons in MCD and control animals. **(A)** Percent of electroporated cells in the cortex of MCD and control mice that express for the neuronal marker NeuN. **(B**,**C)** Neocortical tissue within the EP region immunostained with neuronal marker NeuN at selected time points from P2 to P21 from control (B) and MCD (C) mice. In merged images, magenta represents EP and green represents NeuN. Arrowhead: EP+ and NeuN+, asterisk: EP+ and NeuN-, double arrowhead: EP-and NeuN+. Scale bars 25 μm. ***p = 0.0001. Orientation of all images is such that pial surface is at the top.

### Early upregulation of mTOR expression in EP cells

PS6 is a commonly utilized marker of mTORC1 upregulation: mTORC1 activates the S6Kinases, which in turn phosphorylate S6 (Fig. 1A)^76^. PS6 upregulation has been shown in adult mice in the RhebCA MCD model previously^67^. Presumably, somatic mutations in patients with MCD lead to mTOR pathway upregulation early in development^42^. Thus, to determine whether mTOR pathway upregulation also occurs early during development in the Rheb^CA^ MCD model, we qualitatively examined the expression of phosphorylated ribosomal protein S6 (PS6) via immunohistochemistry at six developmental time points ranging from E18 to P21. We found that there is a relative increase in the intensity of PS6 staining in electroporated neurons vs. the majority of neighboring cells at all examined time points (E18, P2, P7, P14, and P21; Fig. 3: arrowheads). Note that the electroporated cells have larger somata than neighboring cells in the MCD but not control tissue (Fig. 3), which is consistent with the known effect of mTOR upregulation on somatic growth, and is present in human MCDs and animal models of MCD. To our surprise, however, we also noted a small percentage of non-electroporated neurons which demonstrated increased PS6 staining relative to the majority of non-electroporated cells (Fig. 3: double arrowheads). We hypothesize that this surprising finding could be due to mTOR pathway upregulation caused by seizures. Prior studies have shown upregulation of mTOR in models of non-MTOR related epilepsy such as mesial temporal lobe epilespy^77^. Alternatively, it is possible that the “non-EP” cells that exhibit upregulated PS6 received the Rheb^CA^ plasmid but not the GFP plasmid.

**Figure 3.**
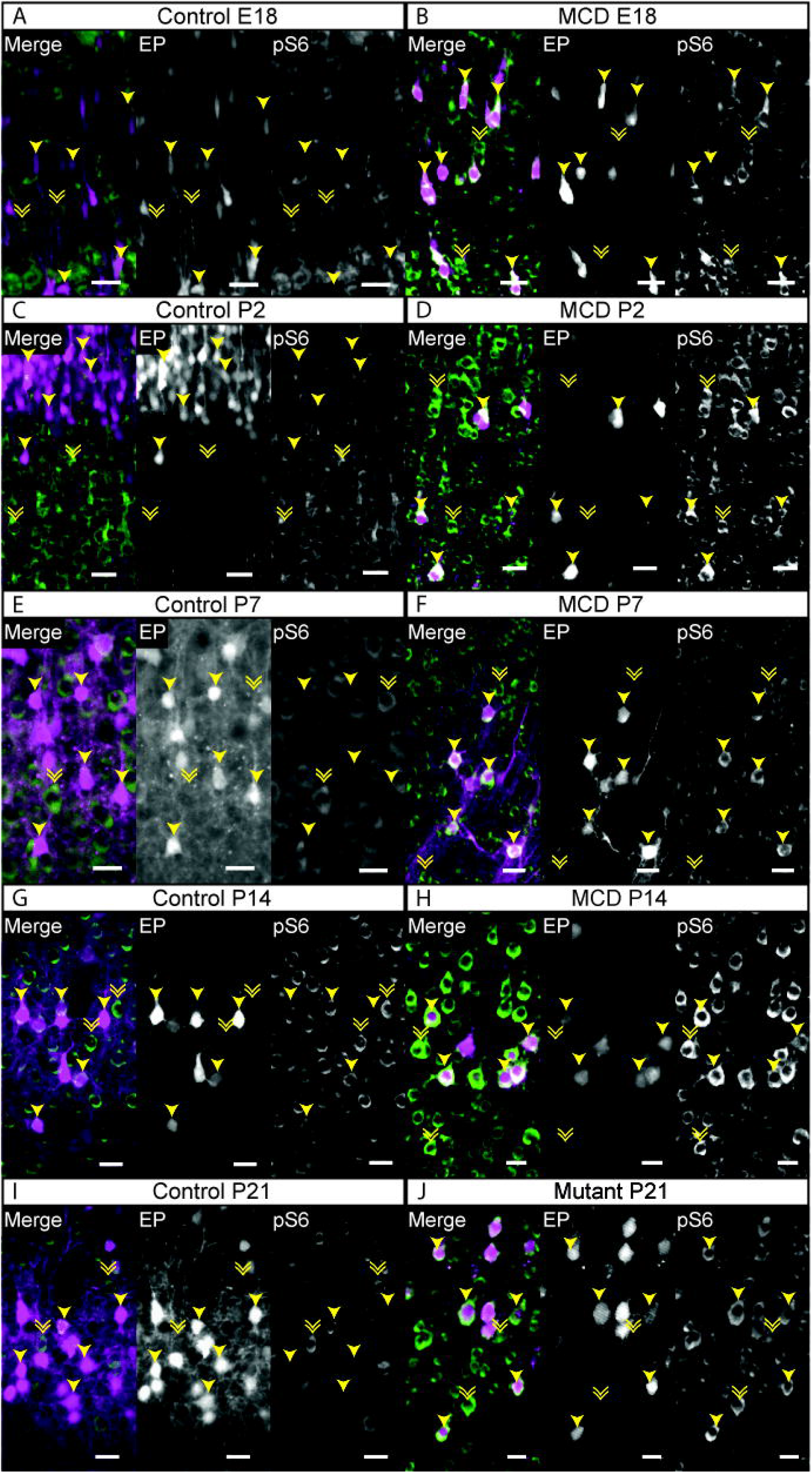
Immunohistochemistry (IHC) of cortical tissue stained for phosphorylated S6 demonstrates upregulation of mTOR in electroporated mice. **(A**,**C**,**E**,**G**,**I):** Control mouse cortical tissue immunostained for PS6 at E18, P2, P7, P14, and P21. **(B**,**D**,**F**,**H**,**J):** Control mouse cortical tissue immunostained for PS6 at E18, P2, P7, P14, and P21. In merged images, magenta represents EP and green represents PS6. Differing brightness between neighboring neurons indicates variation in pS6 expression within the electroporated region. Arrowhead: EP+ and pS6+, double arrowhead: EP- and pS6+. Scale bars: 25 μm. Orientation of all images is such that pial surface is at the top.

### Early differences in neuronal migration patterns

A key feature of focal cortical dysplasia and other malformations of cortical development is dyslamination: i.e. neurons migrate improperly to the cortical layers. In the Rheb^CA^ model of MCD, neuronal precursor cells are electroporated at 15.5. The resulting daughter neurons should be targeted to layer II/III due to the inside-out nature of cortical neuron development. Given that the laminae can be difficult to delineate in MCDs, and that cortical thickness can vary between animals, we chose to express the resultant position of the EP cells in terms of “bins”: each bin represents 1/10^th^ of the distance between the pia and white matter (Fig. 4A). For the bins, layer II/III represents Bins 2-4. At E18, the distribution of cells along the pia-WM axis was similar between control and MCD animals (Fig. 4B,G,L). However, starting at P2, the distribution of neurons for MCD animals was significantly different, with more neurons remaining in the deeper layers (i.e. higher bin numbers). Thus, a large portion of electroporated neurons in the MCD animals do not migrate properly (Fig. 4C-F, H-K, M-P). We would therefore postulate that somatic mutations also occur very early in human development in patients with FCD and HMG to result in very similar histopathological changes.

**Figure 4.**
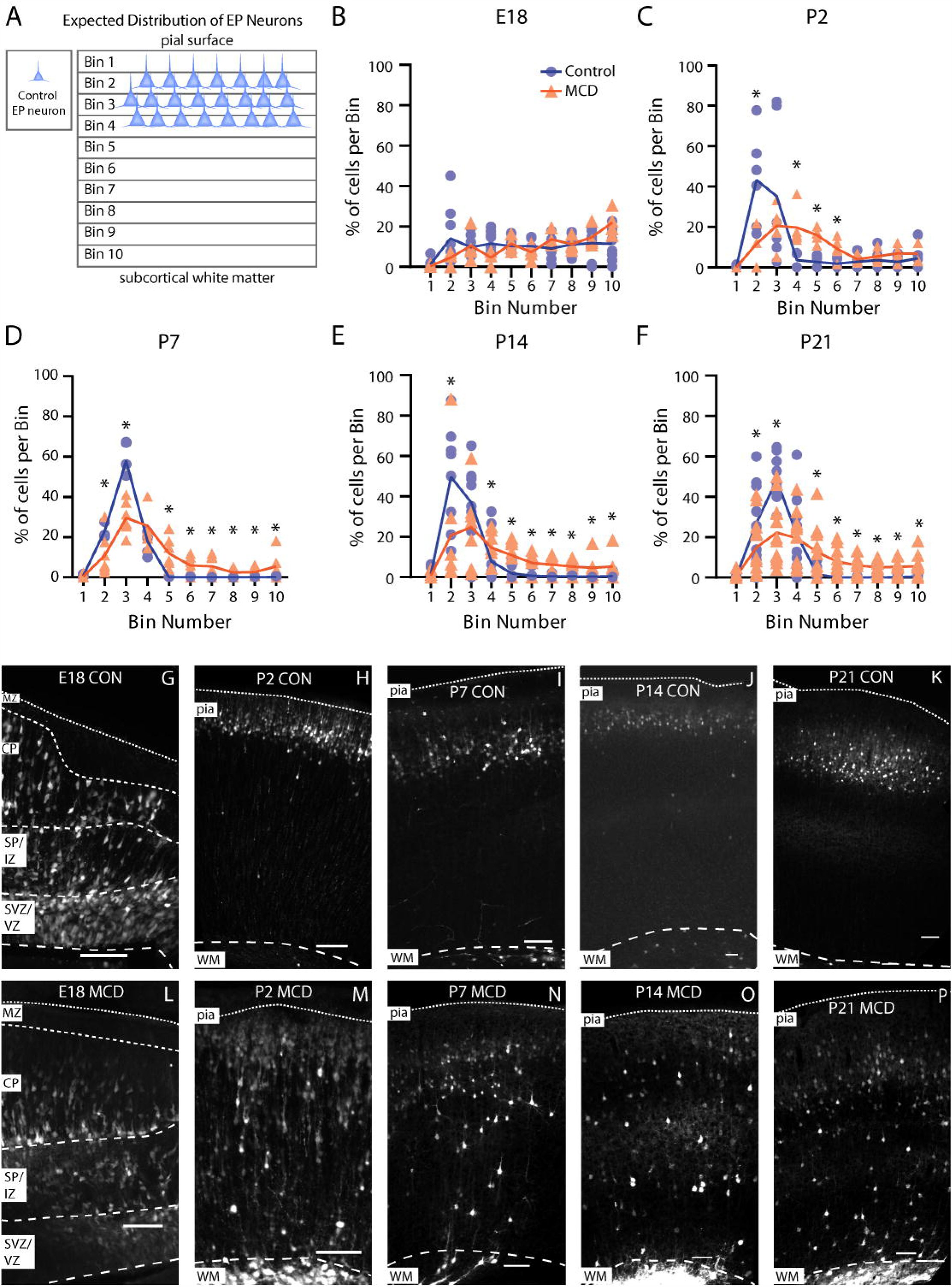
Distribution of electroporated neurons in cortical tissue of electroporated MCD mice indicates a disruption in migration during development. **(A)** Diagram of expected neuronal migration in control mice. Bins 1-10 replaced layers 1-6 for a better quantification of cell location in the cortex with layer 2/3 being represented collectively by bins 2-4. **(B-F)** Frequency distributions of electroporated cells within each bin of the cortex at interval time points through development. **(G-K)** Fluorescent imaging of electroporated neurons in control mice from E18 to P21. **(L-P)** Fluorescent imaging of electroporated neurons in MCD mice from E18 to P21. **(G-P)** At E18, dashed lines indicate marginal zone (MZ), cortical plate (CP), cortical subplate/intermediate zone (SP/IZ), subventricular zone/ventricular zone (SVZ/VZ). At P2-P21, dashed lines indicate pial surface (pia) and white matter border (WM). Scale bars: 50 μm. Significant differences indicated by * in B-F: p < 0.05, FDR = 5%, via multiple Mann-Whitney u tests, using the two-stage step-up method for false discovery rates to correct for multiple comparison.

### Increased axonal arborization in MCD animals

MTOR signaling is known to lead to increases in axon growth. Prior studies in mouse models of MCD show that Rheb overexpression lead to axon length *in vitro* and *in vivo*^*67,78*^. However, we were surprised by the robustness of the axon proliferation that we noted in our animals (Fig. 5J). Given that increased mTOR signaling occurred early in development (Fig. 3), and as migration abnormalies also occurred early (Fig. 4), we postulated that axon overgrowth would similarly occur early in development. We did not detect any increased axon growth at E18 (Fig 5A,B). We found that MCD mice demonstrated large contralateral axon projections from EP neurons starting at P2, and continuing at P7, P14, and P21 (Fig 5C-J).

**Figure 5.**
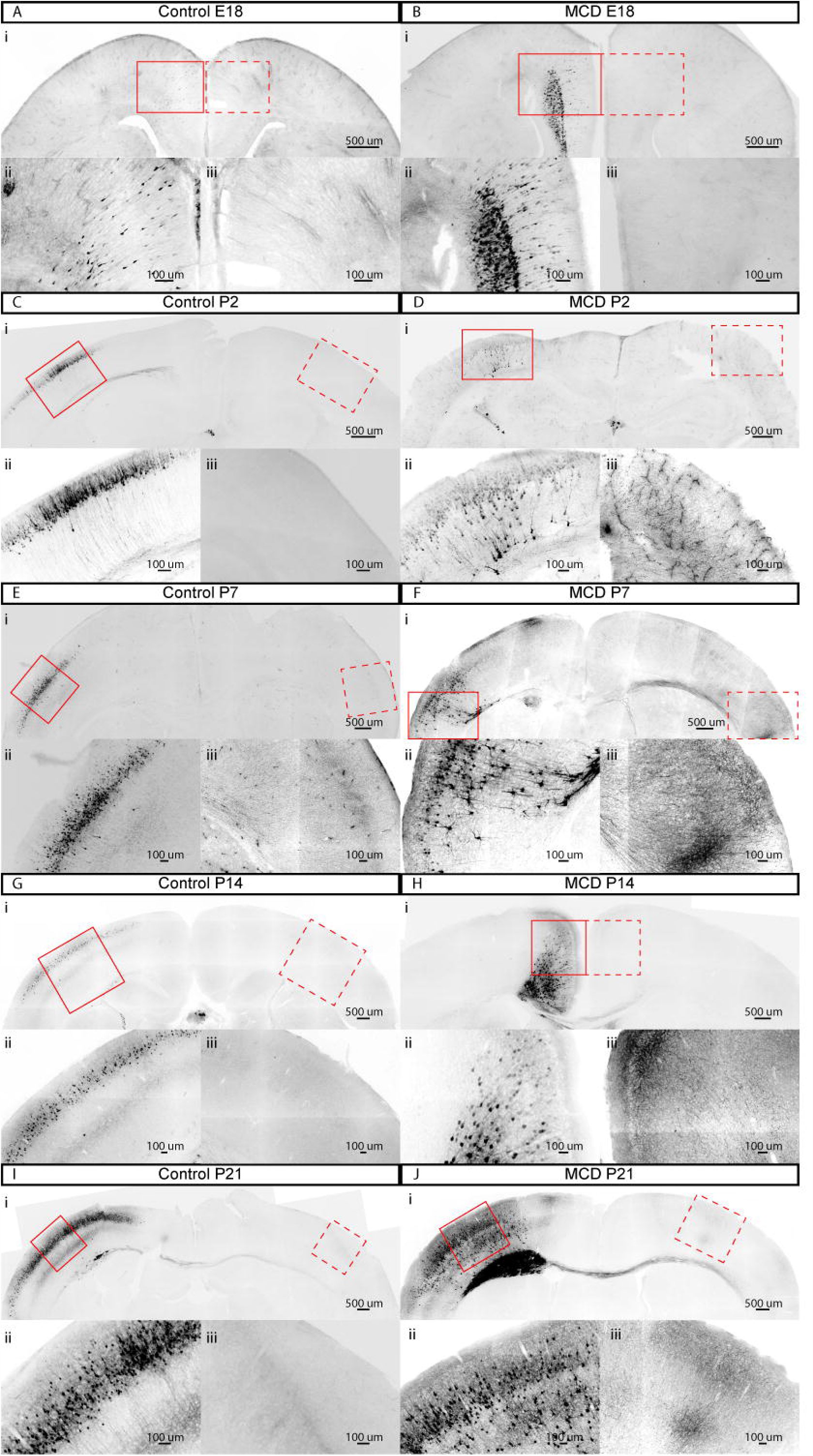
Increased presence of contralateral axonal projections in MCD mice. Fluorescent imaging of dorsal neocortex with magnified views of electroporated region and contralateral homologous region in control and MCD mice at E18 (**A**,**B**), P2 **(C-D)**, P7 **(E-F)**, P14 **(G-H)**, and P21 **(I-J). For A-J:** (i) dorsal brain in coronal section. Solid rectangle indicates electroporated region enlarged in (ii), dashed rectangle indicates contralateral region enlarged in (iii). Note: images are inverted, brighter fluorescence is correlated with darker appearance. Additionally, intensity is adjusted in each panel: in (i) panels, the intensity is balanced to visualize ipsilateral and contralateral fibers, (ii) panels the image intensity is turned down to avoid over-saturation, and in (iii) panels the image intensity is turned up to visualize fine axonal fibers. Scale bars in (i) panels: 500 μm, in (ii) and (iii) panels, 100 μm

## DISCUSSION

In this study, we have demonstrated that, in our hands, Rheb^CA^ mice demonstrate spontaneous recurrent tonic-clonic seizures as well as histological features of MCD. These results support prior data suggesting that *in utero* electroporation of mTOR pathway gain-of-function mutations leads to a histopathologically accurate model of the human condition^44,46,47,67,79-81^. We showed that the effects of mTOR pathway upregulation occur within a few days after electroporation including increased PS6 staining, defects in neuronal migration and lamination, and increased axon length. To our knowledge, this is the first study evaluating the histological changes in mTOR-associated MCDs along a time course of mouse neurodevelopment.

### Axon growth in mTOR-related MCDs

MTOR is a major regulator of transcription and translation, promotes cellular growth, and regulates intracellular autophagy^26-28^. Prior work in both rodent models and cell culture has demonstrated that activation of the mTOR pathway contributes to axonal proliferation during normal development, axonal regeneration after injury, and axonal sprouting after status epilepticus^67,82-91^. Additionally, the transcriptome in axonal growth cones demonstrates a significant enrichment of mTOR-dependent transcripts^92^. Additionally, TSC1/2 loss induces cultured neurons to sprout multiple axons^82^, suppression of PTEN or TSC1 leads to axon regeneration after optic nerve injury^83^, mTOR pathway activation leads to increased numbers of axons in human retinal ganglion cells *in vitro*^*93*^, and mTOR-dependent pathways can contribute to axon repair after spinal cord injury^86,88,94^. Thus, it is not surprising that electroporated neurons in our mouse MCD model demonstrate robust increases in axon growth. This proliferation occurs as early as postnatal day 2.

Our results show that axons show increased callosal projection and contralateral growth in EP neurons which were destined to become layer II/III pyramidal cells. However, with the current experiments, it was not possible to quantify the increase in number or length of axons in an unbiased manner. The number of electroporated neurons in each animal, and the size of the electroporated area itself, as well as the exact location of the electroporated area in the neocortex, Therefore, future studies must be planned to quantify axonal arborization ipsilaterally and contralaterally. We hypothesize that these large increases in axons could contribute to seizure generation by creating a hyper-connected network.

### Mysterious identity of Non-Neuronal EP cells

We were surprised to find that up to 15% of electroporated cells did not express NeuN in the control animals, whereas only 4-7% of MCD neurons were NeuN-negative. We postulated that the increased mTOR signaling in neuronal precursor cells in the EP in MCD animals leads to faster neuronal differentiation in these cells, which would result in fewer cells remaining as glial cells in this developmental pathway. One prior study has suggested that mTOR upregulation leads to faster differentiation in neural stem cells^95^. However, an alternative study suggests that PI3K/AKT signaling (which is upstream of the Rheb mutation induced in our animals), that affects differentiation^96^. Future studies to determine the cellular identify of the EP+, NeuN-cells may help determine the reason for this unexpected finding.

### Non-cell-autonomous upregulation of mTOR pathway

Another surprising finding was that we observed many non-EP neurons in the MCD animals that nevertheless demonstrated increased PS6 immunofluorescence as compared to neighboring cells. Given that mTOR signaling is an intracellular signaling pathway, the simplest explanation for this finding would be that there is not a 100% overlap between the neurons which received the XFP plasmid and the Rheb^CA^ plasmid. However, the vast majority of neurons should indeed express both plasmids. Therefore, we postulate an additional source of mTOR pathway activation other than genetic pathway disruption: seizure-induced activation of this cascade. MTOR dysregulation has been linked to seizure activity in models of epilepsy not based on genetic changes in the mTOR cascade: suppression of mTOR activity with rapamycin prevents axon sprouting and reduces ictal activity in a hippocampal slice culture model of temporal lobe epilepsy^97^, rapamycin treatment can inhibit mossy fiber sprouting (an epileptogenic^98^ increase in aberrant axon connections between hippocampal neurons) in a mouse model of temporal lobe epilepsy^99^, and prevention of axon growth in a mouse model of FCD model by co-electroporation of tetanus toxin leads to seizure suppression^78^. We hypothesize that the seizures that occur in these animals lead to additional mTOR upregulation. Should this hypothesis prove true, this may offer a clue as to why mTOR-related epilepsy remains so difficult to treat: a single mTOR inhibitor may not be sufficient to counteract the powerful upregulation of mTOR that occurs through both genetic and seizure-induced mechanisms. Indeed, there may be parallels between the genetics and treatment of TSC and the genetics and treatment of cancers. Many malignant neoplasms exhibit a “two-hit” pattern within an oncogenic signaling pathway, and treatment of these aggressive lesions requires combination therapy. Similarly, in the pathological lesions of mTOR-related MCDs, successful treatment of these highly epileptogenic lesions may require a combination treatment to attack the mTOR signaling pathway from multiple fronts, especially if there are multiple sources of mTOR upregulation. Future studies are required to unlock the mechanisms behind non-cell-autonomous upregulation of the mTOR pathway in this mouse model, and to determine whether translational implications exist.

## ACKNOWLEDGEMENTS

We would like to thank the EEG core (Michael Mesches) at the University of Colorado Anschutz Medical Campus for their assistance with our studies.

## FUNDING

American Epilepsy Society Junior Investigator Grant to AA.

## Notes

### Competing Interest Statement

The authors have declared no competing interest.

